# Synonymous SNP: Rare versus frequent codon can cause Phenotypic changes in the human genome

**DOI:** 10.1101/582213

**Authors:** Ashu Srivastav

## Abstract

Since the initial sequencing of the human genome, many projects are underway to understand the effects of genetic variation and phenotypic changes between individuals. Single nucleotide polymorphisms (SNPs) are an increasingly important tool for genetic and biomedical research. Synonymous Single Nucleotide Polymorphisms (sSNP) is an important source of human genome variability. It does not produce altered coding sequences therefore expected not to change the function of protein in which they occur. Examination of synonymous SNPs that change a rarely used codon into a frequently used one or vice versa may help in predicting their phenotypic effect on the individual carrying the change. Detail information of Human Synonymous Single-nucleotide-polymorphism may accelerate the research of personalized medicine since it has the crucial impact in the field of non-synonymous SNP.

## INTRODUCTION

Single nucleotide polymorphisms (SNPs) are the most common form of genetic variation in humans comprising nearly 1/1,000th of the average human genome (Taillon-Miller et al. 1998). Traditionally, SNPs are assumed to be biallelic, *i.e.* only two of the four common nucleotides are found in that position, with lower frequency allele being present in at least 1% of the population (Sean Mooney 2005). The distribution and function of SNPs are important areas of current research. We know that synonymous codons (those encoding the same amino acid) are not equally used. This phenomenon is called codon usage bias, which exists in a wide range of biological systems from prokaryotes to eukaryotes. Synonymous codons usage patterns vary significantly among genomes, even often among genes within a single genome. Analysis of codon usage patterns can provide a basis for understanding the mechanism of biased usage of synonymous codons and selecting appropriate host expression systems to improve the expression of target genes in vivo and in vitro (Powell and Moriyama 1997; Zheng et al. 2007).

The variation of synonymous codon preference is correlated with many factors, which are different in organisms. In unicellular organisms, such as *Escherichia coli, Bacillus subtilis, Saccharomyces cerevisiae*, and *Dictyostelium discoideum*, the codon usage is attributable to the equilibrium between natural selection and base compositional mutation bias (Sharp et al. 1993; Shields et al. 1987; Sharp and Li 1986). Reviews are available for understanding how SNPs affect protein structure, the use of SNPs in genetics studies and identifying functional variants in candidate genes. (Marnellos 2004; Rebbeck et al. 2004) Research suggests that most SNPs fall in the 95 per cent non-coding region of the genome (Hagmann 1999).

NCBI’s dbSNP currently contains 4□540□241 validated human entries. Although only 1–3% of the human genome is taken up by protein-coding regions, this small subset of coding SNPs (together with SNPs in gene regulatory regions) has the highest likelihood of being functionally relevant.

The vast majority of prokaryotic and eukaryotic species have non-random codon usage. The major factor in codon choice in many unicellular and some multicellular organisms is Darwinian selection between synonymous codons; highly expressed genes using a restricted set of codons (Gouy and Gautier 1982; Ikemura 1985; Sharp and Matassi 1994) and this selection is almost certainly for optimal translational efficiency, and is most pronounced in highly expressed genes in species whose effective population size is large (Bulmer 1991; Li 1987).

Studies on synonymous codon usage provide information about the molecular evolution of individual genes and can offer data to train methods that detect protein coding regions in uncharacterized genomic DNA (Cancilla et al. 1995b; Freirepicos et al. 1994; Gharbia et al. 1995). Knowledge of synonymous codon usage patterns can also be utilized to design DNA primers and to detect horizontal transfer events (Delorme *et al…* 1994; Groisman et al. 1992; Medigue et al. 1991). Codon-usage analyses can provide insights into the functional categories and histories of genes in a genome. More information can be gained from a codon-usage analysis than a G+C analysis, and it does not require the identification of homologous proteins from other genomes, as is the case for inferring molecular phylogenies. Recent studies have revealed that the heterogeneity in nucleotide composition and the patterns of codon usage bias within many microbial genomes are far more complex than previously imagined (Sharp and Li 1986b; Bulmer 1991). It is well established that synonymous codon usage in various organisms, particularly in unicellular ones, often reflect a balance between the genomic G+C-bias and translational selection (Sharp and Li 1986; Bulmer 1991). The strength and direction of these selection forces vary at the intra and inter-genomic level, greater is the intragenomic bias stronger is the selection for translation efficiency (Musto et al. 2003).

Selection can arise for translational elongation speed or translational accuracy. In *Drosophila*, highly conserved positions more often used preferred codons for amino acids than nonconserved positions in the same gene, suggesting the importance of accuracy (Akashi 1994). Ikemura (1981) showed that abundant proteins of *Escherichia coli* use synonymous codons corresponding to the anticodons of abundant tRNA species. Experimentally, it was found that the correlation is stronger at higher growth rates in *E. coli* in a way that optimizes translation rate (Dong et al. 1996). In the last few years, analyses on the emerging resources of several microbial genomes allowed the perception of several other trends influencing the codon usage. For instance, selection for biased codon usage has also been related with thermophily (Lynn et al. 2002) and the metabolic cost of amino acids (Akashi and Gojobori 2002).

## EXPERIMENTAL SECTIONS

### Tools & Software

E<u>nsembl</u> is a genome browser to visualize and analyze human and many other species genomes. The URL for ensembl genome browser: https://asia.ensembl.org/index.html. Ensembl were used to enlist the information related to sSNP. <u>BioMart</u> is a query-oriented data management and integration system. The URL for biomart tool: http://asia.ensembl.org/biomart/martview/15524ff404a10c7f886a9071124cf6c7. Using Biomart tool were used to retrieve the gene sequence. Python is an easy to learn, powerful programming language. It has efficient high-level data structures and a simple but effective approach to object-oriented programming. Python programming language were used to run individual gene sequences and get the codon list. Codon bias table were used to categories the coding sequence as rare versus frequent codon.

### Synonymous SNP’s data collection

To collect the data related with the synonymous SNP we used the ensembl genome browser. Using the ENSEMBL genome browser with the help of Biomart all the synonymous SNPs in the human genome were obtained. Individual chromosomes were used to retrieve the sSNP by selecting following categories such as: Variant id, Chromosome name, Position on chromosome, Variant alleles, Phenotypic description, Associated gene, Phenotype name, Consequence to transcript, Consequence specific alleles, Transcript strand, Ensembl transcript id, Ensembl gene id, Variation start in CDS(in bps).

### Sequence retrieval

The sequences of each genes containing the sSNP were retrieved from the ENSEMBL genome browser with the help of the Biomart tool by using ENSEMBL gene ID’s. The following parameters were selected: Chromosome name, Ensembl gene id, Coding sequence to retrieve the gene.

### Python programming for finding all the codon of each sequence

Individual gene sequences corresponding to the ensembl gene ID were further processed to get each codon (3 bp’s). To retrieve the codons we used the programming language ‘Python’ to identify all the codons affected by the synonymous SNP in each of the genes. The Human codon bias table were used to select the synonymous codon. The script were written by considering each bp position of the codon. For each run time there is a need to enter the chromosome filename as a raw input. FASTA format of the sequence were used and each individual sequence were recognized by the symbol ‘>’. It will pass the sequence after the run. Three file were generated as “change in first base of codon”, c1, “;second base of codon”, c2,and “third base of codon”, c3 after the run.

### Preparing the list of rare vs frequent codon using the human codon usage table

The human codon usage table were used to enlist the frequency of each codon. Using this table the rare and frequent codons were marked. The criterion used was that a codon is marked as rare if its usage in humans is less than its expected frequency if uniform usage is assumed. For example there are 4 synonymous codons for Ala, so the expected frequency of each codon is 0.25. From the codon usage table it is clear that GCC is a frequently used codon while GCG is a rare codon. Using the above definition of rare and frequent codons the table were prepared.

### Selection of rare vs frequent codon

As per the list of rare and frequent codons, codons for both the alleles were marked manually as rare or frequent codon.

## RESULTS

### Synonymous snp’s data collection

The result shows that all the selected parameters were obtained in a tabulated form in excel sheet. The respective column shows the related information which is necessary to process further.

**Table: 1.**
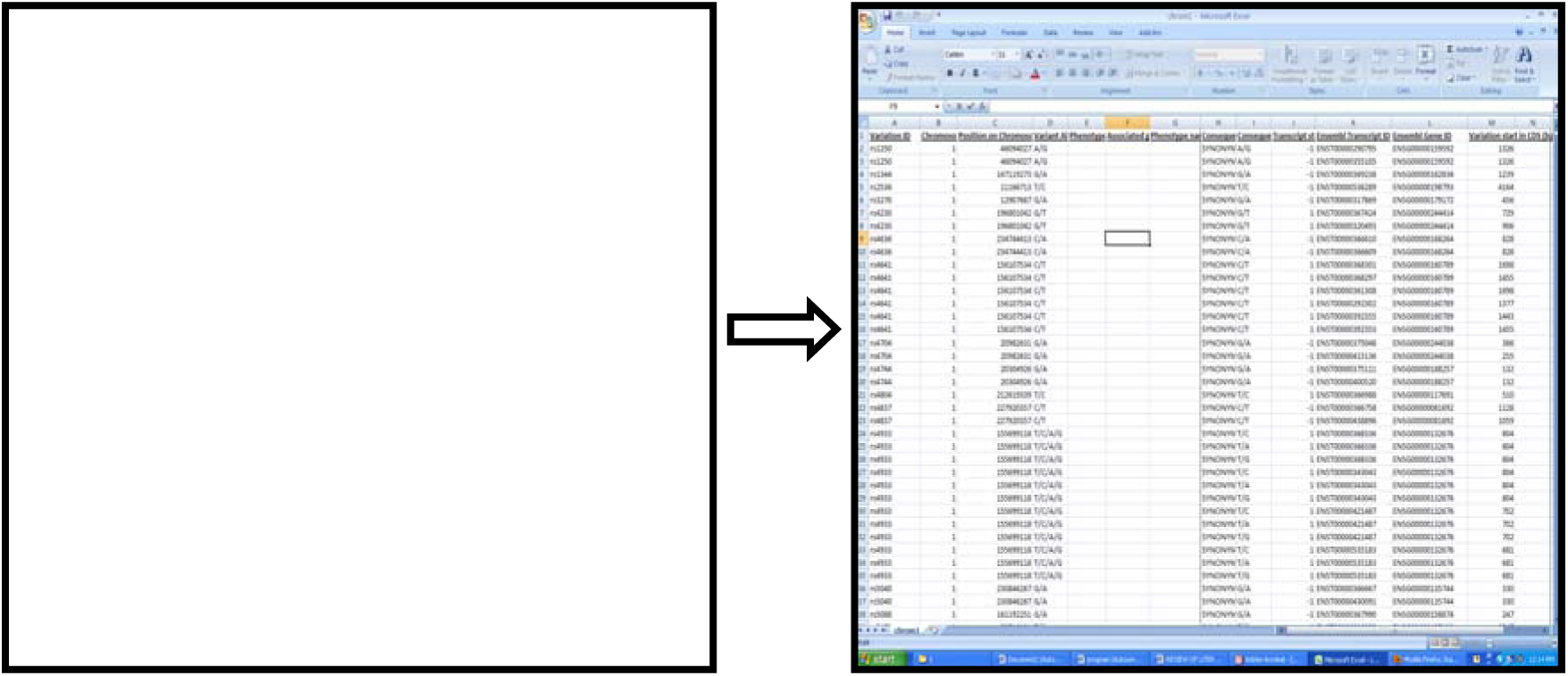
a. Ensembl genome browser, b. Retrieving the selected parameters of sSNP data.

### Sequence retrival

Biomart tool were used to get the home sapiens gene which gives the sequence in FASTA format. Each generated notepad shows a series of genes sequences in FASTA.

**Table: 2.**
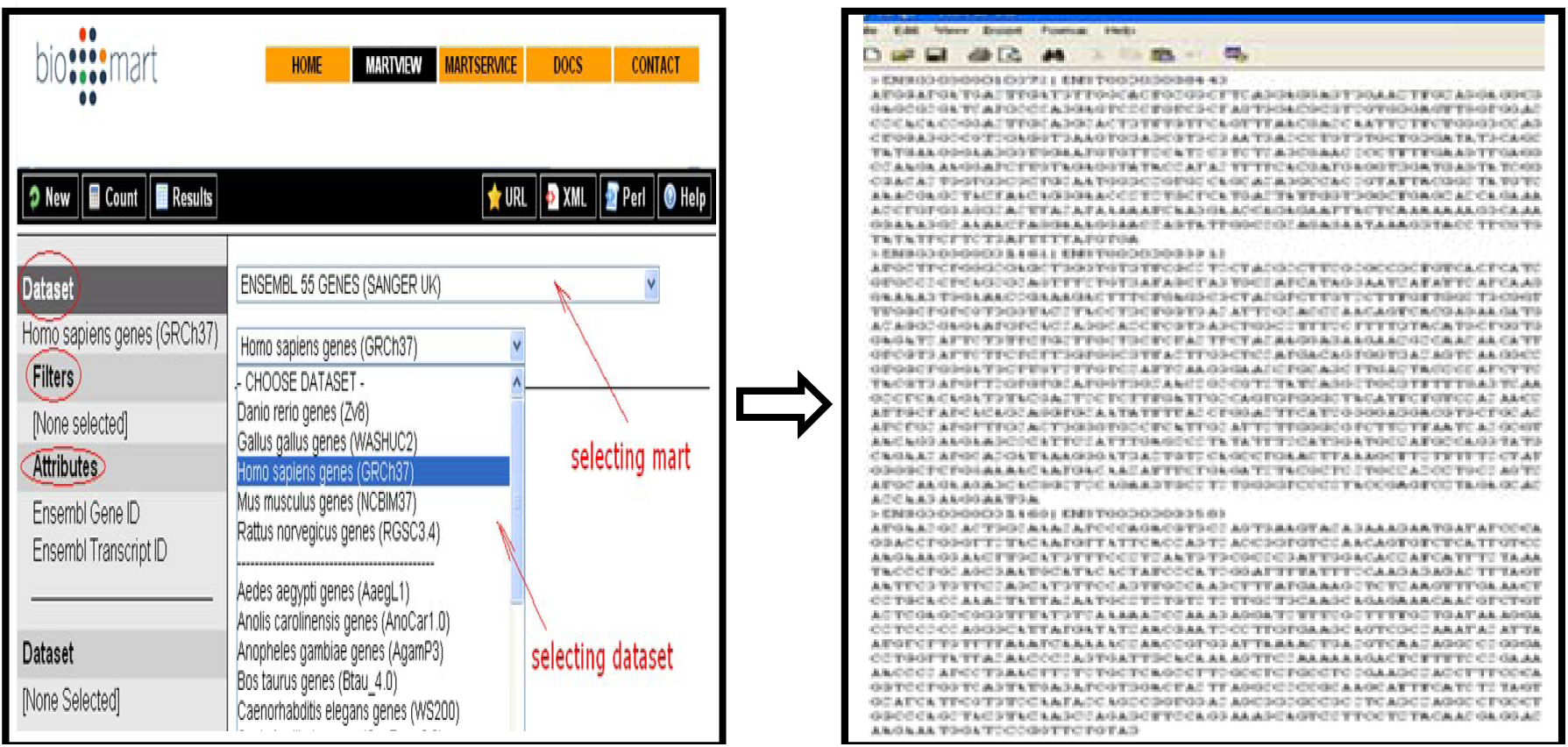
a. Biomart tool selecting parameters, b. Extraction of gene sequences using Biomart.

### Synonymous codon list

As a output of the compiled python program, a separate file for each codon position (1,2 or 3) affected by the SNP were generated. All three different files will be named according to their Ensembl gene id. The excel file shows the ensembl gene id, ensembl transcript id and the coding amino acid name, transcript strand as plus or minus strand. The synonymous codon of each variant were also enlisted as triplet bases.

**Table: 3.**
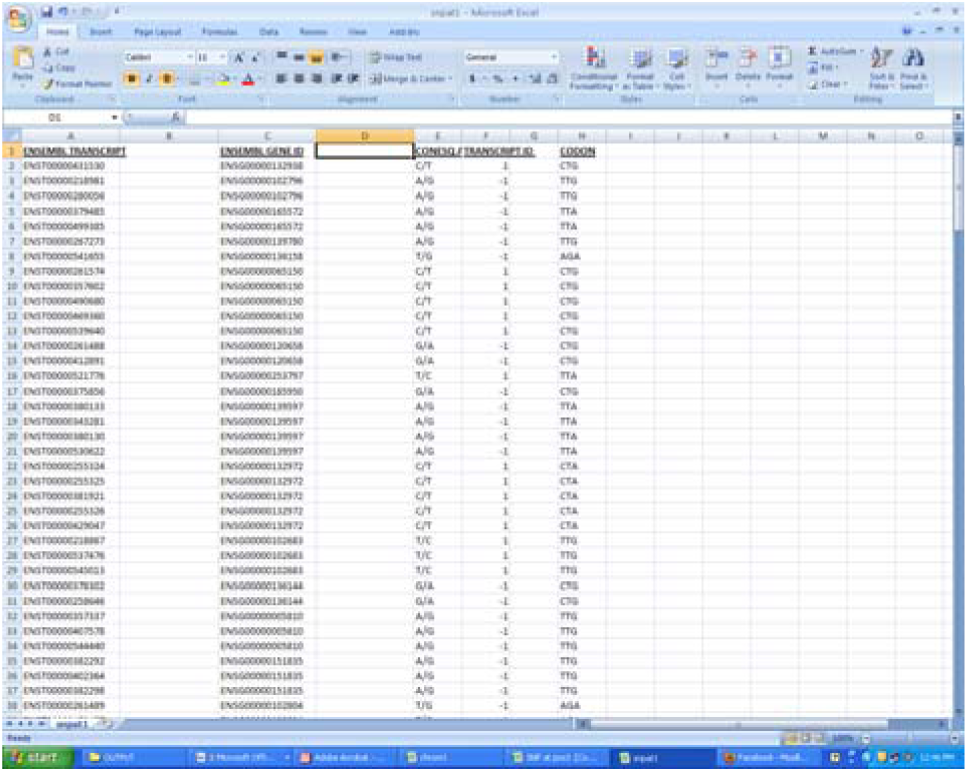
Codon listing of each Ensembl gene ID.

### Rare or frequent codon list

The prepared list of codon frequency (rare *vs.* frequent codon) were used to categorised the ensembl gene sequences. Each synonymous codons were categorised as previous codon frequency with the changed one. The changed codon frequency were enlisted as rare to frequent or vice-versa based on their allelic code. All the data were prepared and stored for further analysis associated with this codon frequency changes.

**Table: 4.**
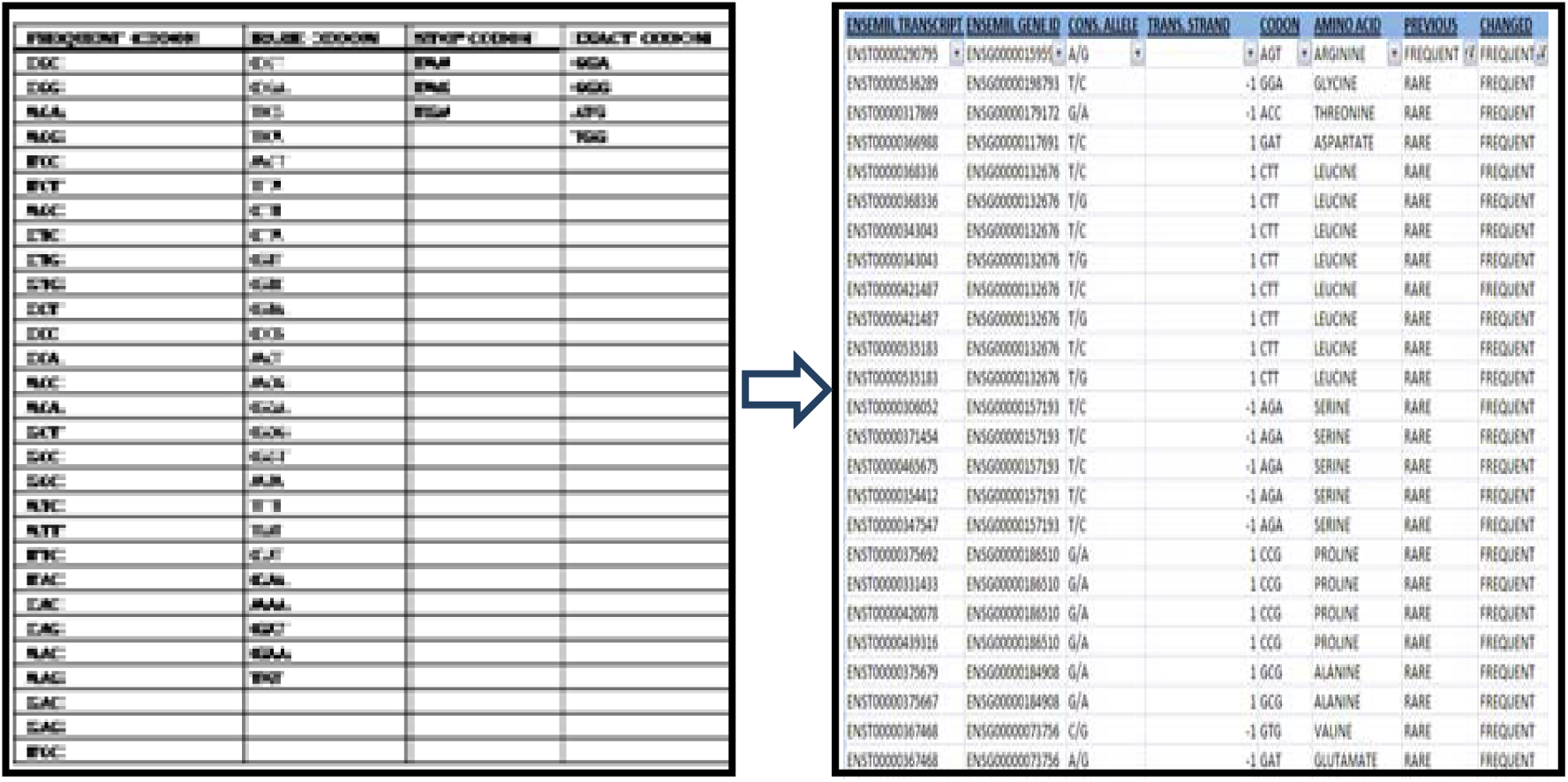
a. Codon listing as per their frequency, b. Assigning the codon frequency changes as per the allele.

## DISCUSSIONS

Synonymous SNP which don’t change the amino acid sequence plays a very crucial role in personalized medicine. Synonymous variation can have effects on potential patho-physiological and pharmacogenetics. Studies shows that the presence of a rare codon, as sSNP, affects the timing of co-translational folding and even the insertion of P-gp into the membrane, resulting the alteration in the structure of substrate and inhibitor interaction sites. Even it was also observed that in some cases synonymous changes in humans have been shown to cause genetic disorders by exon skipping. While in the case of GC content at third position of codons (GC3) classically used to investigate whether natural selection influences synonymous codon usage the departure between the GC3 of a gene and the GC frequency observed in adjacent non-coding regions which can be used as a measure of gene specific forces (e.g. selection) on synonymous mutations beyond isochoric mutational tendencies. Those genes showing the strongest departure from isochoric GC content are the best candidates for the detection of functional changes resulting from synonymous mutations. In case of protein folding it has been observed that the synonymous codon usage can affect the conformational state of the protein. It also shows that codon specific translation rate may influence the in-vivo protein folding. 3 dimensional structure of the protein were affected by the rare codon as the presence of rare codon in turns, loops and domains linkers has been observed. Further studies are required to find out more phenotypic changes caused by synonymous SNP’s.

## ACKNOWLEDGEMENTS

I would like to thank Mr. Ankit Agarwal for helping me in coding language, Dr. Raja G. and Dr. Manish Kesharwani for their contribution in literature searching and necessary dealing with files. The work was performed under the guidance of Dr. Shrish Tiwari, Centre for Cellular and Molecular Biology and successfully completed.

